# Honeybee dance-followers respond similarly to dances regardless of their spatial information content

**DOI:** 10.1101/2021.03.24.436796

**Authors:** Matthew J. Hasenjager, William Hoppitt, Ellouise Leadbeater

## Abstract

Honeybees famously use waggle dances to communicate foraging locations to nestmates in the hive, thereby recruiting them to those sites. The decision to dance is governed by rules that, when operating collectively, are assumed to direct foragers to the most profitable locations with little input from potential recruits, who are presumed to respond similarly to any dance regardless of its information content. Yet variation in receiver responses can qualitatively alter collective outcomes. Here, we use network-based diffusion analysis to compare the collective influence of dance information during recruitment to feeders at different distances. We further assess how any such effects might be achieved at the individual level by dance-followers either persisting with known sites when novel targets are distant and/or seeking more accurate spatial information to guide long-distance searches. Contrary to predictions, we found no evidence that dance-followers’ responses depended on target distance. While dance information was always key to feeder discovery, its importance did not vary with feeder distance, and bees were in fact quicker to abandon previously rewarding sites for distant alternatives. These findings provide empirical support for the longstanding assumption that self-organized foraging by honeybee colonies relies heavily on signal performance rules with limited input from recipients.

## Introduction

Living in groups provides opportunities to pool information across multiple individuals in order to make accurate collective decisions (e.g. navigation in homing pigeons [1]; predator avoidance in fish [2]). In the social insects, such decisions are the product of many (often thousands of) individual-level environmental assessments that are shared with nestmates through evolved communication signals. Simple rules that govern the production or longevity of these signals can generate non-linear feedbacks that produce accurate collective decisions [3-5]. A classic example involves waggle dance-based recruitment to foraging locations in the western honeybee (*Apis mellifera*), whereby energetically efficient trips elicit more waggle runs on return to the hive [6,7]. Closer sites should hence be over-represented on the dancefloor, and thus attract more recruits, relative to distant alternatives that offer resources of similar quality. This straightforward performance rule could thus enable colonies to collectively optimize energetic efficiency without requiring that dance-followers use the spatial information contained in the dance to make any decision about the potential value of the trip that lies ahead of them [3].

In the above scenario, dance-followers are expected to respond similarly to any dance, regardless of its content. Yet research over the past decade has revealed the sophisticated ways in which insects acquire, process, store, retrieve, and use information [8], raising the possibility that signal recipients decide how to respond by weighing the costs and benefits of using that information. For example, ants generally ignore trail pheromones in favour of memories, but will switch to trail-following if information indicates that doing so will lead to a higher quality food source [9]. Likewise, experienced honeybee foragers often discount the spatial information contained in dances in favour of returning to known foraging locations [10–12] and may devalue dance information when it repeatedly proves unreliable [13]. Accounting for such individual variation in receiver responses can lead to qualitatively different outcomes in models of collective behaviour [3,14,15].

Here, we use network-based diffusion analysis (NBDA; [16,17]) to evaluate the responses of dance-followers to dances that indicate novel close or distant feeders. NBDA can provide an estimate (*s*) of the influence of each dance circuit followed on a dance-follower, and we propose that this influence may decrease with distance to the target when dance-followers are unfamiliar with the target resource. This is because locating new sites can require multiple search trips and hence significant time costs that potentially increase with distance [6,18]. We created pools of unemployed yet motivated foragers and allowed their recruitment to either close or distant feeders, estimating the strength of social transmission through the resulting dance networks. We further monitored behaviour at the individual level to establish the mechanisms by which such collective effects might be achieved, predicting that (i) bees that follow dances for distant target recruitment sites may persist with known sites for longer, rather than attempting to locate the new food source, and that (ii) the same bees may invest in gaining more accurate location information by following more waggle runs pre-departure [19,20]. Finally, we monitored individual dancer behaviour to confirm our expectation, based on previous work [6,7], that closer resources will be over-represented on the dancefloor.

## Methods

### Colony housing

These experiments were carried out on the campus of Royal Holloway, University of London from July – September 2018. Three queen-right honeybee colonies were housed indoors within three-frame observation hives with unrestricted access via tunnels to the outdoors. Colonies contained 2000 – 3000 workers, brood, and reserves of pollen and honey. Each colony underwent both a short-distance and a long-distance recruitment trial, performed consecutively to minimise differences in colony and environmental conditions across trials (Table S1).

### Training

Working with a single colony at a time, two groups of foragers (13 – 31 per group) were simultaneously trained using standard techniques (described in [21,22]) to two feeders providing unscented sucrose solution. In each case, one feeder was designated the *recruit feeder* (always 100m from the hive) and the other the *target feeder* (either 100m or 500m from the hive) with an angular separation of ~110° between the two feeders (figure 1). During training, foragers were assigned unique enamel paint marks upon first arriving at a feeder, meaning we could be confident that individuals trained to the recruit feeder had never visited the target feeder. Later, during the test period (see below), the recruit feeder would become depleted, creating a pool of marked potential recruits for the target feeder (figure 1).

**Figure 1.**
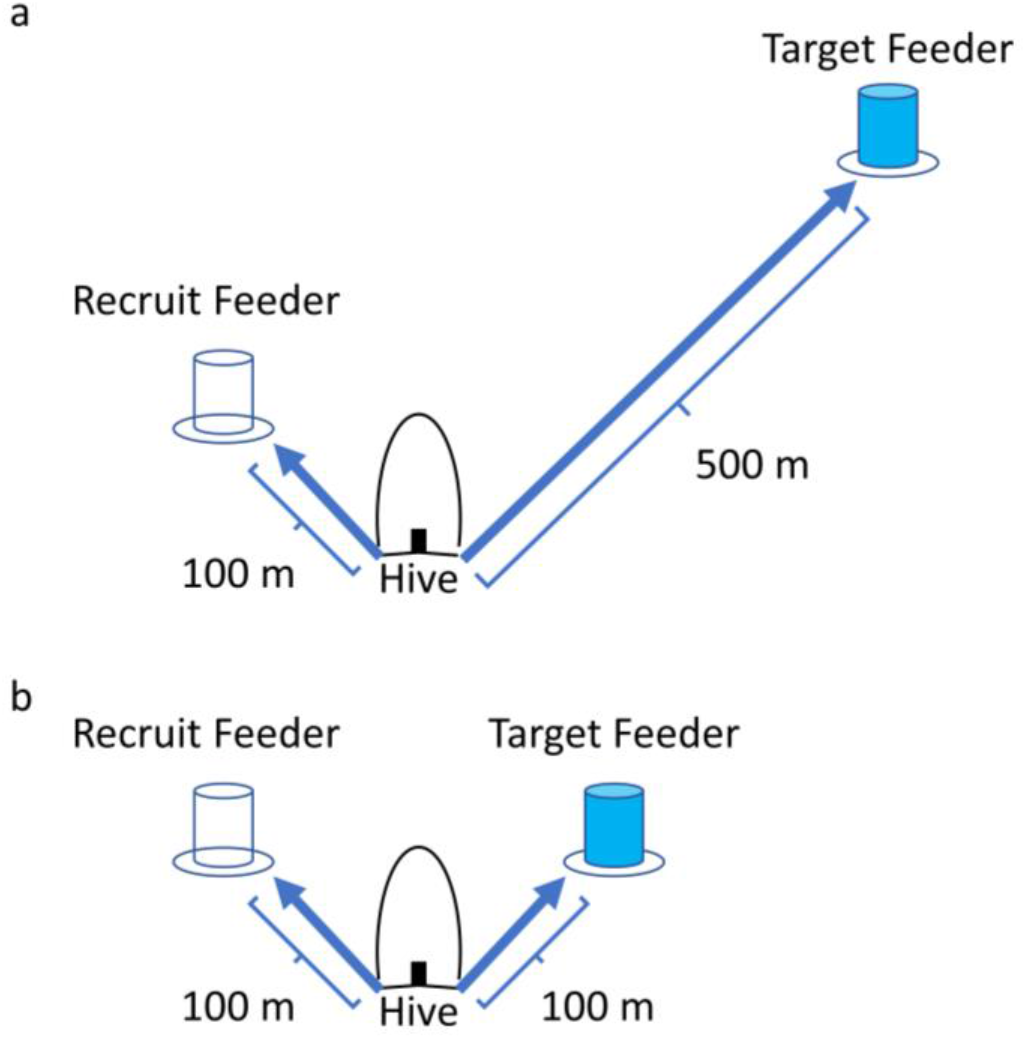
Feeder arrangement used during (a) long-distance and (b) short-distance trials. Within each trial, cohorts were simultaneously trained to the recruit and target feeders. During trials, the recruit feeder was left empty to create a pool of potential recruits for the target feeder.

Training took place over 5 – 11 days per trial. Both feeders offered identically scented sucrose on the final day of training for one hour (following [22]: 50 μL essential oil per L sucrose, plus reservoir of essential oil below the feeder; scents varied between trials; table S1) in order to promote greater interest in the target feeder during the trial [see 11]. Only individuals that visited the recruit feeder during either this odour presentation or during the previous training day were used as potential recruits during the trials. Although most potential recruits visited the recruit feeder multiple times during both of these training days, a small number of individuals only visited the feeder once during the odour presentation or were only observed during one 30 min census on the previous day. However, as excluding these 21 individuals did not qualitatively alter our findings or conclusions, we retained them in the full analysis.

### Trials

Trials commenced between 0930 – 1000 on the following morning. During a trial, the target feeder continued to provide scented 2M sucrose, whereas the recruit feeder was left empty (figure 1), thus mimicking a common natural scenario in which one tree or patch of flowers comes into bloom at the same as another of the same species ceases to be rewarding. We allowed 10 – 12 foragers previously trained to the target feeder to collect from it, while any remaining members of this cohort were captured upon arrival. Successful recruits from the recruit feeder were also allowed to collect freely from the target feeder. We did not restrict the activities of other bees in the hive, but any that located the target feeder were captured on arrival. Using both video recordings and in-person observations, we recorded arrival and departure times for each marked individual at both the recruit and target feeders throughout the trial. Trials lasted either 120 min (colony A) or 180 min (colonies B and C); this change was implemented to allow recruits in the 500 m trials additional time to locate the target feeder.

During trials, we filmed the dance floor within the observation hive. A wooden baffle directed foragers onto one side of the hive, meaning the vast majority of dances were visible. For each hive visit made by target feeder foragers (including recruits), we recorded its duration, whether dancing occurred, and the number of waggle runs produced. We also recorded all dance-following interactions between marked individuals, noting participant identities, when each interaction occurred, its duration (sec), and the number of waggle runs followed. A bee was defined as following a waggle run if its head was oriented towards the dancer within 1 antennal length [12]. We further recorded the occurrence of waggle dances by other bees in the hive for natural food sources, whether these dancers carried pollen, the number of waggle runs produced, and any instances in which a marked individual followed one of these dances.

### Network-based diffusion analysis (NBDA)

All analyses were carried out in R ver. 4.0.3 [23]. In an NBDA, the strength of social transmission per unit of network connection (e.g. per waggle run followed), relative to individual exploration, is estimated by the social transmission parameter, *s* [16,17]. Here, we set out to compare estimates of *s* between the close and distant feeders, based on social networks constructed from our video records of dance-following interactions. Specifically, we used order-of-acquisition diffusion analysis, in which networks are used to predict the order in which individuals acquire a behaviour—here, discovery of the target feeder within each trial [17]. Network connections were directed from dancers to followers, and we included models where connections were weighted either by the number of waggle runs followed or the total duration (sec) of dance-following in our candidate model sets (described below). To capture the temporal ordering of dance-following interactions, we used dynamic networks that updated when individuals departed the hive for the target feeder [24].

To compare the relative influence of dance-based transmission for recruitment across our distance treatments, we fit models in which *s* was either estimated separately for short- and long-distance trials (*s_short_* ≠ *s_Long_*) or in which *s* was constrained to be equal across these treatments (*s_short_* = *s_Long_*). See the Supplementary Material for more details on specification of the NBDA models and for the complete candidate model set. Due to asymmetry in the uncertainty for parameter estimates, profile likelihood techniques were used to obtain 95% CIs [25]. The NBDA was carried out using the *NBDA* package [26].

### Individual-level analyses

Prior to seeking out a new feeder, honeybee foragers typically return to known sites (often extremely persistently), even if they know those sites to be unrewarding [27]. To examine potential differences in this persistence, and in pre-departure information gathering, when the alternative target feeder is either close or distant, we classified trips where individuals were observed at the recruit feeder as “reactivation” trips. If instead that recruit left the hive for more than 90 seconds and successfully discovered the target feeder or was not observed at either site, it was classified as searching for the target feeder (‘search trip’).

A full description of the individual-level analyses, including all fixed and random effects in each global model, is provided in tables S2 and S3 and summarised here. Our primary analyses focussed on the effects of target distance on follower behaviour in terms of: (i) the number of waggle runs followed before departing the hive (zero-inflated negative binomial GLMM); and (ii) the probability of searching for the target feeder vs reactivating during these absences (binomial GLMM). To confirm that longer target distances incur greater search costs, we also analysed (iii) the duration of hive absences (linear mixed-effects model); and (iv) the number of unsuccessful searches prior to locating the target feeder (Poisson GLMM).

For completeness, we also analysed dancer behaviour across the short- and long-distance treatments, to compare how the two target feeders were represented on the dancefloor. We included (i) hive visit frequency (linear mixed-effects model); (ii) mean hive visit duration (linear mixed-effects model); (iii) the probability of dancing per visit (binomial GLMM); and (iv) the mean number of waggle runs produced during visits with dancing (linear mixed-effects model).

In every model, *Trial* and *colony* were included as a random intercept term and fixed effect respectively; *individual* was included as a random effect for analyses that included multiple observations per individual. All input variables were mean-centred and continuous variables were scaled by dividing by twice their standard deviation [28,29]. LMMs were fitted using nlme [30] to model heteroscedasticity in the residuals [31] and GLMMs were fitted with glmmTMB [32]. Inspection of GLMM residuals was carried out using DHARMa [33].

We performed model selection on all candidate models nested within each global model (tables S2 and S3) on the basis of AICc. Models were removed from the candidate set if they were more complex versions of a model with a lower AICc value [29,34,35]. From this reduced model set, we extracted a 95% confidence set of models and used these to obtain model-averaged parameter estimates (MAEs), unconditional standard errors (USEs), and unconditional 95% confidence intervals (CIs) [34]. Where a single model received especially strong support (*w_i_*, ≥ 0.95), inferences were based on this model alone. Multimodel inference was performed using the MuMIn package [36].

## Results

### Network-based diffusion analysis (NBDA)

In the short- and long-distance trials respectively, 49 and 25 recruits successfully located the target feeder (table 1). Dance information was key in guiding to foragers to the target feeder, regardless of its distance from the hive. Of our candidate set for the NBDA, two models received nearly all support (model probabilities: *w*_1_ = 0.91; *w*_2_ = 0.09). Both included the dance-following network and constrained social transmission rates to be equal across distance treatments (i.e. *s*_100 *m*_ = *s*_500 *m*_), indicating that the acceleratory effects of dance-based transmission over how rapidly individuals discovered the target feeder did not vary with foraging distance. The two models differed only in how network connections were weighted: the top-ranked model weighted connections according to the number of waggle runs followed, whereas the second-ranked model used the total duration of dance-following interactions. The best-supported model estimated a social transmission rate of 2.42 x 10^7^ (95% CI: 0.90, +∞), corresponding to an estimated 97 – 100% of recruitment events explained by dance-following. Estimates from the second-ranked model yielded essentially identical results. In summary, the NBDA indicated that successful recruitment was predicted by an individual’s investment in dance-following but provided no evidence that the influence of dance information differed according to the indicated distance. See table S4 for parameter estimates from both models.

**Table 1.** Summary of experimental trials (TF: target feeder; RF: recruit feeder). Data provided as sample size or mean ± SD. Hive absences were labelled as ‘reactivation’ if a forager returned to the RF; otherwise, foragers were assumed to be searching for the TF.

| Colony | TF<br>distance | RF<br>trained | TF<br>recruits | Dancers: Waggle<br>runs per hive visit | Followers: TF waggle runs<br>followed per hive visit |  |
| --- | --- | --- | --- | --- | --- | --- |
|  |  |  |  |  | Reactivation | Searching |
| A | 100 m | 21 | 16 | 11.4 ± 18.5 | 3.9 ± 4.2 | 7.4 ± 5.3 |
| A | 500 m | 26 | 3 | 11.8 ± 15.3 | 2.3 ± 3.5 | 8.3 ± 6.4 |
| B | 100 m | 31 | 22 | 15.5 ± 15.5 | 5.2 ± 6.9 | 14.4 ± 7.9 |
| B | 500 m | 22 | 9 | 12.5 ± 13.6 | 3.6 ± 4.6 | 17.4 ± 10.2 |
| C | 100 m | 28 | 11 | 4.6 ± 8.1 | 2.3 ± 3.3 | 4.7 ± 4.8 |
| C | 500 m | 30 | 13 | 11.3 ± 13.6 | 4.1 ± 7.1 | 13.6 ± 8.5 |

### Follower behaviour

As expected, individuals typically made multiple trips to the empty recruit feeder before searching for the new target (see also [11]), and the probability of abandoning the recruit feeder in favour of searching for the target feeder increased over time (binomial GLMM: hive visit: MAE ± USE = 2.7 ± 0.23 (95% CI: 2.25, 3.15); figure 2). However, contrary to our expectations, bees were quicker to engage in search trips when the target feeder was distantly located than when it was close to the hive (binomial GLMM: target feeder distance (500 m): MAE ± USE = 0.9 ± 0.22 (95% CI: 0.48, 1.32); target feeder distance * hive visit: MAE ± USE = 1.91 ± 0.46 (95% CI: 1.01, 2.81); figure 2). See tables S5 and S6 for full model summaries.

**Figure 2.**
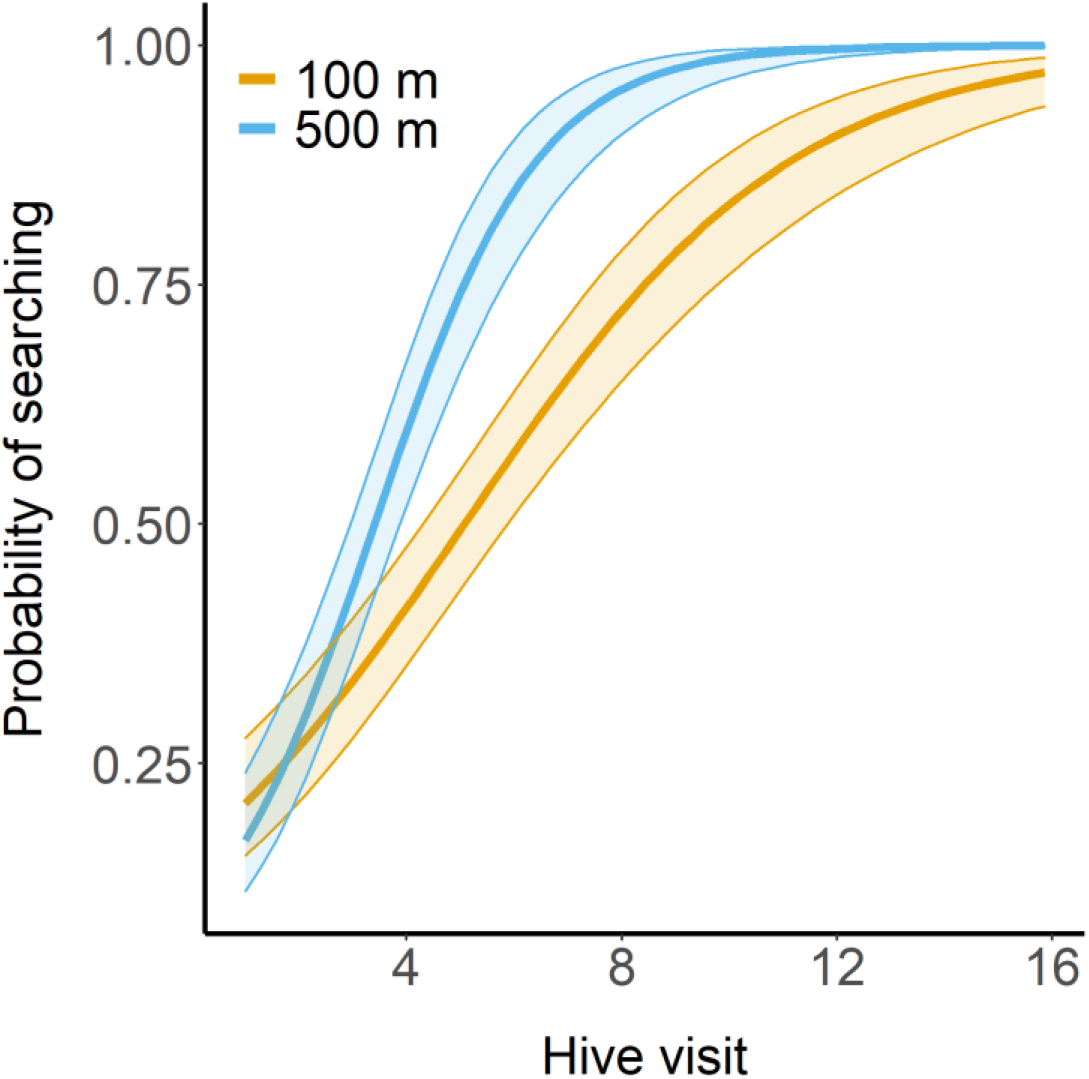
Predicted probability of searching for the target feeder upon departing the hive. Lines and shaded regions respectively indicate predicted values and 95% CI calculated from model-averaged GLMM fixed effects with all random effects set to 0.

In line with previous work [11,37], we found that foragers on average followed more waggle runs before departing in search of the target feeder than when re-visiting the empty feeder, though this effect lessened over time (zero-inflated negative binomial GLMM: search trip: MAE ± USE = 0.75 ± 0.06 (95% CI: 0.63, 0.87); searching * hive visit: MAE ± USE = −0.53 ± 0.13 (95% CI: −0.78, −0.27)). However, there was no evidence at the 95% confidence level that bees followed more waggle runs before searching when the feeder was more distantly located (same GLMM: TF distance (500 m) * search trip: MAE ± USE = 0.17 ± 0.15 (95% CI: −0.11, 0.47); figure 3). See tables S7 and S8 for full model summaries.

**Figure 3.**
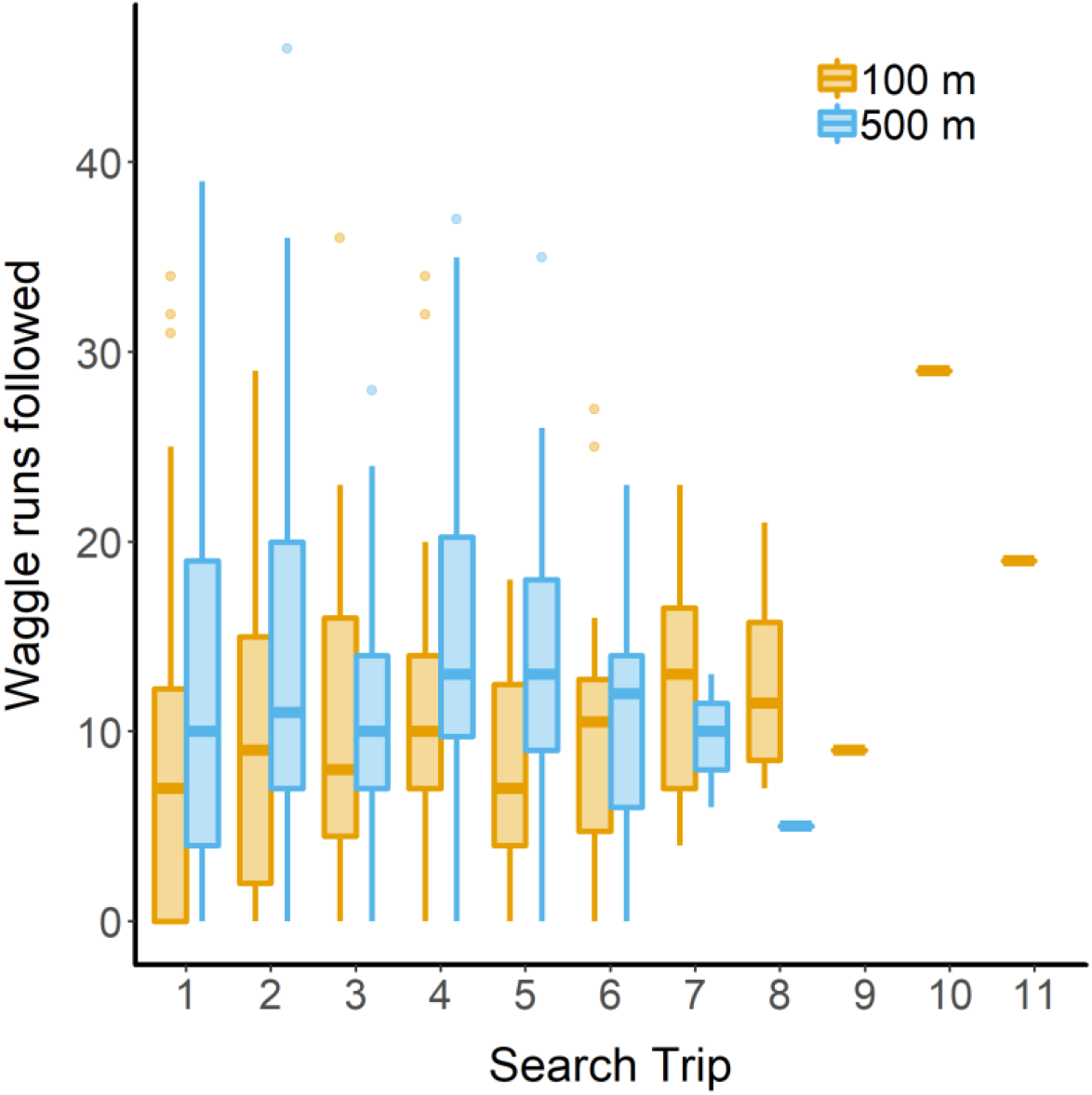
The number of waggle runs followed prior to searching for the target feeder. Thick lines indicate medians, boxes enclose the interquartile range, and whiskers extend to up to 1.5x this range.

As expected if long-distance searches are more costly, search trips into the field were longer in duration than reactivation trips only when the target feeder was more distant (LMM: target feeder distance (500 m) * search trip: MAE ± USE = 0.6 ± 0.06 (95% CI: 0.48, 0.72); figure 4a; tables S9 & S10). Comparing the mean duration of searches for the target feeder vs. collection trips made by employed foragers (minus time spent at the feeder) confirmed that both search and collection trips took more time when the target feeder was more distant from the hive (LMM: target distance (500 m): estimate ± SE = 0.63 ± 0.03 (95% CI: 0.5, 0.76); table S11) and that search trips were longer in duration than collection trips (LMM: trip type (collection): estimate ± SE = −0.6 ± 0.05 (95% CI: −0.69, −0.51); table S11)). However, searches were not disproportionately longer at 500 m than at 100 m (the best-supported model, *w_i_* > 0.99, did not include an interaction between target feeder distance and trip type; table S11). Regardless of distance, successful recruits undertook a similar number of unsuccessful searches before eventually locating the target feeder (Poisson GLMM: target feeder distance (500 m): MAE ± USE = −0.24 ± 0.22 (95% CI: −0.67, 0.18); figure 4b; tables S12 & S13).

**Figure 4.**
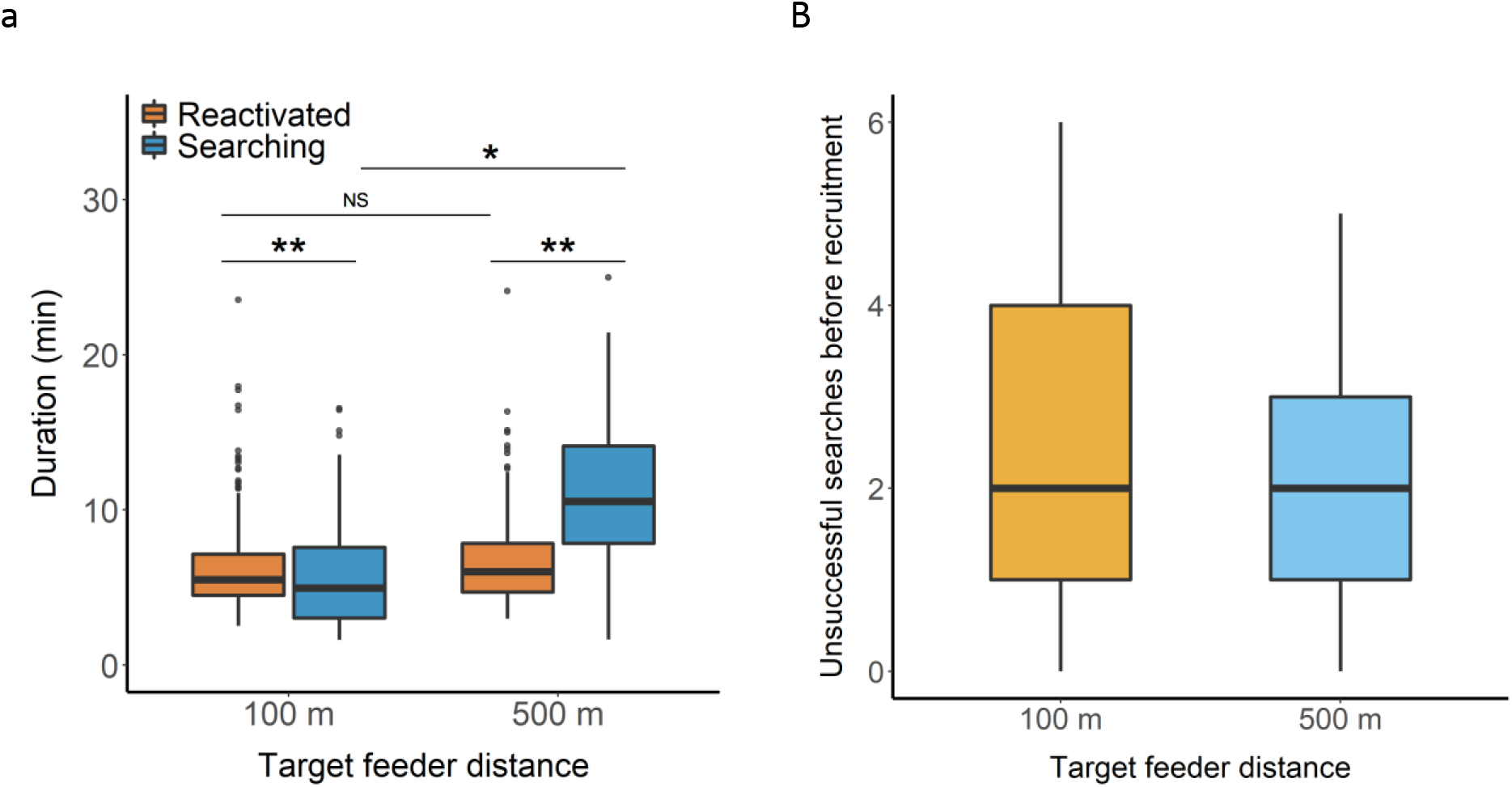
(a) Duration of hive absences and (b) number of unsuccessful searches before discovering the target feeder. Absences were labelled as reactivations if foragers returned to the empty recruit feeder and searches for the target feeder otherwise. The y-axis in (a) has been truncated to enhance clarity; an additional reactivation was observed in both the short- and long-distance trials with respective durations of 64.9 and 35.7 min. Thick lines indicate medians, boxes enclose the interquartile range, and whiskers extend up to 1.5x this range. *P* values for contrasts were adjusted using the Bonferroni method: *: P < 0.01; **: P < 0.001.

In addition to dances for the target feeder, we observed 122 dances for natural food sources. These dancers produced 8.74 ± 12.37 (mean ± SD) waggle runs per dance and carried pollen in 56 of these dances. Although our focal bees occasionally followed these natural dances, these following events were brief in duration (mean ± SD = 1.13 ± 0.35 waggle runs followed; n = 40 dance-following events). Out of 519 prospective search flights, only 10 (i.e. 1.9%) involved a focal bee following a dance for a natural food source (mean ± SD = 1.8 ± 1.03 waggle runs followed).

### Dancer behaviour

In line with previous work [7,21], dances representing the more distant (and thus less energetically efficient) 500 m feeder were underrepresented on the dancefloor relative to those for the closer feeder. This occurred because dancers visited the hive less frequently when the target feeder was more distantly located (LMM: target feeder distance (500 m): estimate ± SE = −3.58 ± 0.37 (95% CI: −5.15, −2.01); table S14). On average, dancers made 10.3 visits hr^-1^ when the feeder was located 100 m from the hive, but only 6.9 visits hr^-1^ when it was 500 m away (figure 5a). This is in part because travel to and from the distant feeder took longer (table S11), but foragers collecting at 500 m also tended to remain in the hive for longer on each visit (LMM: target feeder distance (500 m): estimate ± SE = 0.51 ± 0.09 (95% CI: 0.12, 0.89); figure 5b; table S15). In contrast to our expectation that bees foraging at the distant feeder would be less likely to dance upon returning to the hive, there was no evidence that foraging distance influenced foragers’ propensity to dance (table S16). If anything, foragers in long-distance trials tended to be more likely to dance during hive visits (binomial GLMM: target feeder distance (500 m): estimate ± SE = 0.72 ± 0.71 (95% CI: −0.67, 2.11)), though the best-supported model (*w_i_* > 0.99) did not include this effect (table S16). There was also no evidence that dancers for more distant feeders produced fewer waggle runs (LMM: target feeder distance (500 m): MAE ± USE = 0.31 ± 1.79 (95% CI: −3.2, 3.82); tables S17 & S18; figure 5c).

**Figure 5.**
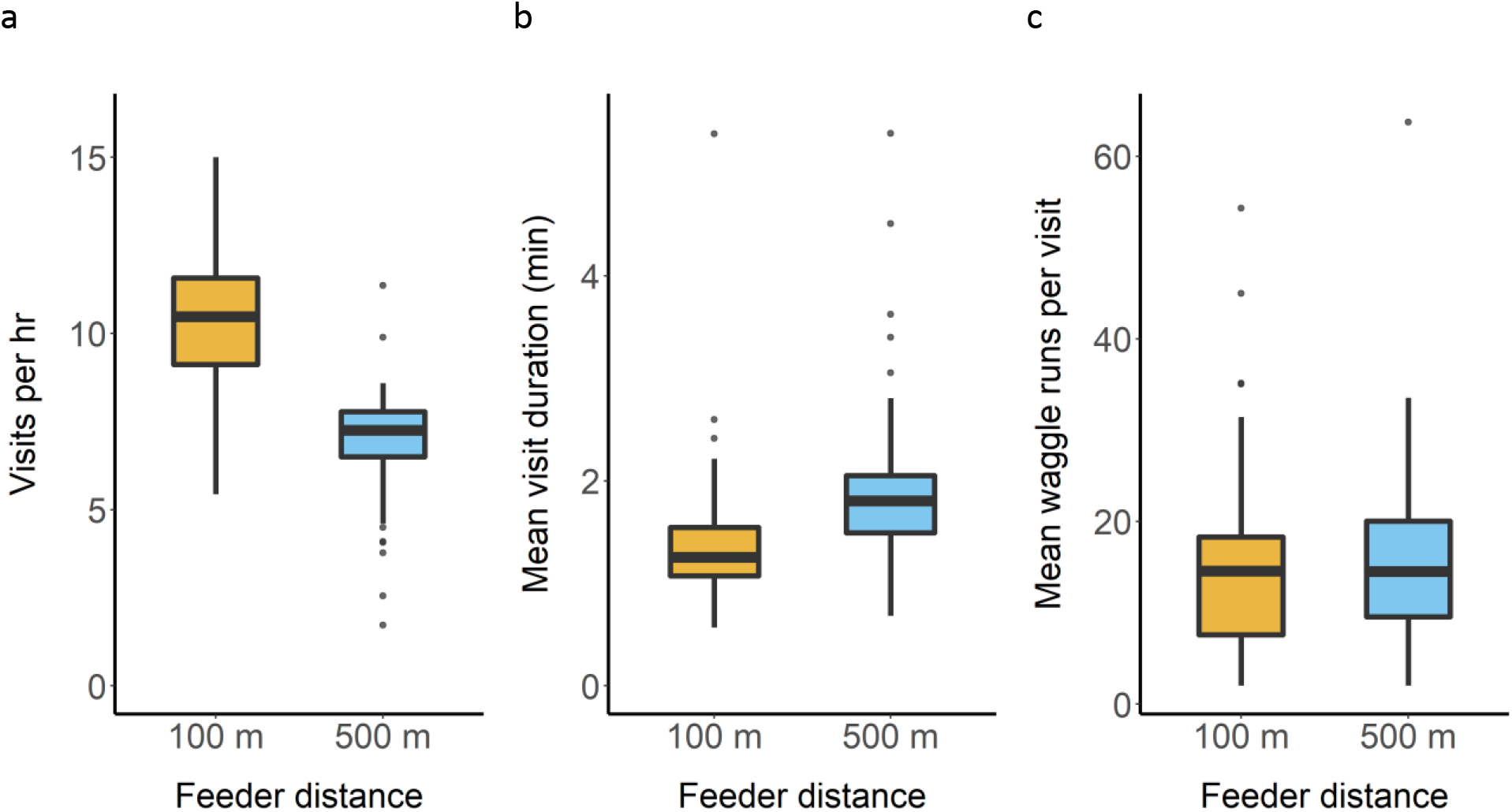
(a) Hive visit frequency, (b) mean visit duration, and (c) mean waggle runs produced per visit with dancing by foragers collecting from the target feeder. Thick lines indicate medians, boxes enclose the interquartile range, and whiskers extend up to 1.5x this range.

## Discussion

The traditional view of insects as mere stimulus-response “machines” has given way to a growing recognition that despite their miniature brains, insects possess sophisticated cognitive capabilities [8]. Thus, although empirically derived theoretical models have shown how simple rules that govern the production of waggle dances are sufficient to generate adaptive collective responses by honeybee colonies without requiring that dance-followers evaluate the transmitted spatial information [3,22], dance-followers may in principle be able to fine-tune their responses according to this information. Here, we used NBDA to first ask whether bees respond differently to dances depending on the indicated foraging distance. In contrast to our predictions, we found no difference in the estimated influence of dance communication (*s*) on the order in which recruits arrived at our close or distant novel feeders. We further found that increased foraging distance was not associated with increased investment in dance-following prior to searching, and that foragers were in fact quicker to abandon a depleted site when the alternative was more distantly located. Taken together, these findings suggest that dance-followers do not evaluate the distance information contained in a dance when deciding how to respond to it.

As foraging distances increase, searches require progressively greater investments in time and energy, exacerbated by the fact that dance-guided searches often fail (figure 4b; [6,18]). Why then are bees not more reticent to accept recruitment to distant novel sources? Dances are followed both by bees that have never visited the target site (recruitment) and bees that know its location (reactivation), with only the former incurring search costs. Since dancers do not know for which purpose their audience is following, we expected the behavioural rules that translate the energetic efficiency of a foraging trip into the number of waggle runs performed to ignore these additional search costs, allowing instead for dance-followers to fine-tune their responses depending on their informational status. However, it may be that the increasing search costs elicited by distant resources are already sufficiently accounted for through their under-representation on the dancefloor and that additional receiver responses are not needed to achieve adaptive collective foraging, especially given that we found that although search costs do increase with distance, they are not disproportionately large at greater distances.

Alternatively, it may be the case that while sensitivity to distance information by followers could increase colony foraging efficiency, the mechanisms by which it could be achieved have diminishing returns. For example, beyond a certain point, the extra time spent following additional dance circuits may not appreciably increase the likelihood of locating a site. Moreover, the positive relationship between foraging site distance and waggle run duration means that as foraging distances increase, foragers must invest ever more time in dance-following to acquire similar amounts of information [21,38]. The use of dance information may involve a speed-accuracy trade-off [39], such that setting out with reasonably accurate spatial information may often be preferable to investing further time in waiting for and following dances.

Although honeybees have been known to forage from sites that are located upwards of 10 km from the hive [22,40], dance decoding studies have shown that the median distance travelled under natural conditions is often an order of magnitude lower than this [41]. For example, we recently found the median distance indicated by dances across an entire season in southern England to be 708m and 1108m for urban and agricultural sites respectively [42]. Nonetheless, it is clear that our feeder locations, at 100 and 500m from the hive, do not represent the full foraging range. However, we note that previous work has detected modifications to dance behaviour between sites at 250 and 500m [7], and that our distance treatments were distinct enough to drive observable differences in search costs. Thus, while we cannot rule out that dance followers take the indicated distance into account when deciding whether to seek out a very distant site, we are confident that our treatments should have elicited an effect if one exists within this range.

In agreement with earlier studies [11,12,37], most foragers visited the empty recruit feeder several times before searching for the target feeder. Yet rather than foragers being more reluctant to abandon this site when alternatives were more distant (as predicted), the opposite pattern was observed (figure 2). It is possible that this finding simply stems from how foraging trips were labelled—i.e., during reactivations, it was assumed that bees did not also search for the target feeder. However, studies using harmonic radar to track bees’ foraging flights have revealed the occurrence of such cross-trips between familiar and unfamiliar foraging locations [43], potentially allowing individuals to gain up-to-date information on familiar foraging sites while also making use of dance information without requiring that they first return to the hive. If joint reactivation-search trips occurred more often in short-distance trials when feeders were relatively close together [43], this could be reflected in our analysis as a lower likelihood of searching when the target was nearby. However, although we cannot rule out that such trips occurred, our data suggest that they were unlikely to be especially common (see Supplementary Material, tables S19 & S20). Alternately, the dance-indicated location in long-distance trials may have been easier to identify as a novel site, as neither the distance nor directional components matched that of the recruit feeder [21,37]. Regardless, our results complement previous reports that honeybees’ persistence to familiar sites depend more on previous profitability than on the availability of alternatives [27].

Although we assumed that during departures from the hive, potential recruits were either returning to the recruit feeder or searching for the target feeder, individuals may also have engaged in alternative foraging behaviours, including visiting other known foraging locations or searching for natural food sources. However, trials took place during the late summer and early autumn when few natural food sources are available to bees in southern England [41]. Accordingly, foragers were highly persistent in visiting the feeders during training, limiting their opportunities to learn about other foraging sites prior to the trial. In addition, there were relatively few dances for natural sources during the trials and these were rarely followed by our focal individuals. When natural dances were followed, these bouts were always brief in duration, indicating that foragers were not attempting to decode the dance’s spatial information [44]. Individuals may also have attempted to locate other foraging sites through individual scouting. However, previous reports have found that scouting is relatively rare when dances are readily available in the hive [18,45], as was the case in our study. We therefore feel confident that most searching events represented attempts to locate the target feeder. Nevertheless, we repeated our analysis of: (i) the duration of searching events and (ii) the number of waggle runs followed prior to each search using only the subset of successful recruitment events. Our findings were consistent with our more inclusive analysis: in long-distance trials, searches were longer in duration and recruits followed more waggle runs prior to a successful search, but this latter difference was not significant at the 95% level (tables S21 & S22).

Given that the colony represents the reproductive unit in honeybees, natural selection is expected to have acted on the heuristics that guide behaviour at the individual level in order to produce adaptive colony-level responses [3]. Although such individual-level algorithms could in principal lead recruits to differentially respond to dances according to the indicated distance, we found no evidence that this is the case. Rather, our results provide empirical support to the long-standing assumption that the effective allocation of recruits among foraging sites does not depend on information processing by dance-followers, but on the rules that govern the production of dances themselves, the tempo of foraging, and whether or not to abandon a foraging patch [22]. However, due to the challenges involved in studying decision-making in bees foraging on natural sources, most studies (including our own) have used artificial food sources located relatively near to the hive that offer an unrestricted flow of sucrose. Additional investigations into how the production of dances is modulated under more naturalistic foraging conditions and how dance-followers respond to this information would be worthwhile.

## Data accessibility

Raw data and code to reproduce all analyses are available from the Dryad Digital Repository: https://doi.org/10.5061/dryad.8kprr4xn8 [46].

## Authors’ contributions

M.J.H. and E.L. designed the study. M.J.H. collected the data and M.J.H. and W.H. analysed it. M.J.H. wrote the initial draft and all authors contributed to revisions.

## Acknowledgements

We thank Alex Hadleigh for assistance with carrying out the experiments described here.

## References

1. Dell’Ariccia, G., Dell’Omo, G., Wolfer, D. P., & Lipp, H.-P. (2008). Flock flying improves pigeons’ homing: GPS track analysis of individual flyers versus small groups. Animal Behaviour, 76, 1165–1172.

2. Ward, A. J. W., Herbert-Read, J. E., Sumpter, D. J. T., & Krause, J. (2011). Fast and accurate decisions through collective vigilance in fish shoals. Proceedings of the National Academy of Sciences of the U. S. A., 108, 2312–2315.

3. Detrain, C., & Deneubourg, J.-L. (2008). Collective decision-making and foraging patterns in ants and honeybees. Advances in Insect Physiology, 35, 123–173.

4. Seeley, T. D. (2010). Honeybee democracy. Princeton, NJ: Princeton University Press.

5. Sasaki, T., Granovskiy, B., Mann, R. P., Sumpter, D. J. T., & Pratt, S. C. (2013). Ant colonies outperform individuals when a sensory discrimination task is difficult but not when it is easy. Proceedings of the National Academy of Sciences of the U. S. A., 110, 13769–13773.

6. Seeley, T. D., & Towne, W. F. (1992). Tactics of dance choice in honey bees: do foragers compare dances? Behavioral Ecology and Sociobiology, 30, 59–69.

7. Seeley, T. D. (1994). Honey bee foragers as sensory units of their colonies. Behavioral Ecology and Sociobiology, 34, 51–62.

8. Giurfa, M. (2015). Learning and cognition in insects. Wiley Interdisciplinary Reviews: Cognitive Science, 6, 383–395.

9. Czaczkes, T. J., Beckwith, J. J., Horsch, A.-L., & Hartig, F. (2019). The multi-dimensional nature of information drives prioritization of private over social information in ants. Proceedings of the Royal Society B, 286, 20191136.

10. Grüter, C., Balbuena, M. S., & Farina, W. M. (2008). Informational conflicts created by the waggle dance. Proceedings of the Royal Society B, 275, 1321–1327.

11. Grüter, C., & Ratnieks, F. L. W. (2011). Honeybee foragers increase the use of waggle dance information when private information becomes unrewarding. Animal Behaviour, 81, 949–954.

12. Hasenjager, M. J., Hoppitt, W., & Leadbeater, E. (2020). Network-based diffusion analysis reveals context-specific dominance of dance communication in foraging honeybees. Nature Communications, 11, 625.

13. l’Anson Price, R., Dulex, N., Vial, N., Vincent, C., & Grüter, C. (2019). Honeybees forage more successfully without the “dance language” in challenging environments. Science Advances, 5, eaat0450.

14. Schürch, R., & Grüter, C. (2014). Dancing bees improve colony foraging success as long-term benefits outweigh short-term costs. PLoS ONE, 9, e104660.

15. Lemanski, N. J., Cook, C. N., Smith, B. H., & Pinter-wollman, N. (2019). A multiscale review of behavioral variation in collective foraging behavior in honey bees. Insects, 10, 370.

16. Franz, M., & Nunn, C. L. (2009). Network-based diffusion analysis: a new method for detecting social learning. Proceedings of the Royal Society B, 276, 1829–1836.

17. Hoppitt, W., Boogert, N. J., & Laland, K. N. (2010). Detecting social transmission in networks. Journal of Theoretical Biology, 263, 544–555.

18. Biesmeijer, J. C., & Seeley, T. D. (2005). The use of waggle dance information by honey bees throughout their foraging careers. Behavioral Ecology and Sociobiology, 59, 133–142.

19. Tanner, D., & Visscher, K. (2008). Do honey bees average directions in the waggle dance to determine a flight direction? Behavioral Ecology and Sociobiology, 62, 1891–1898.

20. Tanner, D., & Visscher, K. (2009). Does the body orientation of waggle dance followers affect the accuracy of recruitment? Apidologie, 40, 55–62.

21. von Frisch, K. (1967). The dance language and orientation of bees. Cambridge, MA: Harvard University Press.

22. Seeley, T. D. (1995). The wisdom of the hive: The social physiology of honey bee colonies. Cambridge, MA: Harvard University Press.

23. R Core Team. (2020). R: a language and environment for statistical computing. R Foundation for Statistical Computing, Vienna, Austria. https://www.R-project.org/

24. Hasenjager, M. J., Leadbeater, E., & Hoppitt, W. (2021). Detecting and quantifying social transmission using network-based diffusion analysis. Journal of Animal Ecology, 90, 8–26. doi: 10.1111/1365-2656-13307.

25. Morgan, B. J. T. (2009). Applied stochastic modelling (2nd ed.) Boca Raton, FL: Chapman & Hall/CRC Press.

26. Hoppitt, W., Photopoulou, T., Hasenjager, M., & Leadbeater, E. (2020). NBDA: a package for implementing network-based diffusion analysis. R package version 0.9.4. https://github.com/whoppitt/NBDA

27. Al Toufailia, H., Grüter, C., & Ratnieks, F. L. W. (2013). Persistence to unrewarding feeding locations by honeybee foragers *(Apis mellifera):*the effects of experience, resource profitability and season. Ethology, 119, 1096–1106.

28. Schielzeth, H. (2010). Simple means to improve the interpretability of regression coefficients. Methods in Ecology and Evolution, 1, 103–113.

29. Grueber, C. E., Nakagawa, S., Laws, R. J., & Jamieson, I. G. (2011). Multimodel inference in ecology and evolution: challenges and solutions. Journal of Evolutionary Biology, 24, 699–711.

30. Pinheiro, J., Bates, D., DebRoy, S., Sarkar, D., & R Core Team. (2018). nlme: linear and nonlinear mixed effects models. R package version 3.1–137, https://CRAN.R-project.org/package=nlme.

31. Zuur, A. F., Ieno, E. N., Walker, N. J., Saveliev, A. A., & Smith, G. M. (2009). Mixed effects models and extensions in ecology with R. New York, NY: Springer Science+Business Media.

32. Brooks, M. E., Kristensen, K., van Benthem, K. J., Magnusson, A., Berg, C. W., Nielsen, A., et al. (2017). glmmTMB balances speed and flexibility among packages for zero-inflated generalized linear mixed modeling. The R Journal, 9, 378–400.

33. Hartig, F. (2020). DHARMa: residual diagnostics for hierarchical (multi-level/mixed) regression models. R package version 0.3.1. https://CRAN.R-project.org/package=DHARMa.

34. Burnham, K. P., & Anderson, D. R. (2002). Model selection and multimodel inference: A practical information-theoretic approach (2nd Ed.). New York, NY: Springer-Verlag.

35. Richards, S. A. (2008). Dealing with overdispersed count data in applied ecology. Journal of Applied Ecology, 45, 218–227.

36. Barton, K. (2019). MuMIn: multi-model inference. R package version 1.43.6. https://CRAN.R-project.org/package=MuMIn.

37. Grüter, C., Segers, F. H. I. D., & Ratnieks, F. L. W. (2013). Social learning strategies in honeybee foragers: do the costs of using private information affect the use of social information? Animal Behaviour, 85, 1443–1449.

38. Al Toufailia, H., Couvillon, M. J., Ratnieks, F. L. W., & Grüter, C. (2013). Honey bee waggle dance communication: signal meaning and signal noise affect dance follower behaviour. Behavioral Ecology and Sociobiology, 67, 549–556.

39. Chittka, L., Skorupski, P., & Raine, N. E. (2009). Speed-accuracy tradeoffs in animal decision making. Trends in Ecology and Evolution, 24, 400–407.

40. Beekman, M., & Ratnieks, F. L. W. (2000). Long-range foraging by the honey-bee, *Apis mellifera* L. Functional Ecology, 14, 490–496.

41. Couvillon, M. J., Schürch, R., & Ratnieks, F. L. W. (2014). Waggle dance distances as integrative indicators of seasonal foraging challenges. PLOS ONE, 9, e93495.

42. Samuelson, A. E., Schürch, R., & Leadbeater, E. (2019). Dancing bees evaluate agricultural forage resources as inferior to central urban land. bioRxiv, 2019.12.19.882076.

43. Menzel, R., Kirbach, A., Haass, W.-D., Fischer, B., Fuchs, J., Koblofsky, M., et al. (2011). A common frame of reference for learned and communicated vectors in honeybee navigation. Current Biology, 21, 645–650.

44. Grüter, C., & Farina, W. M. (2009). The honeybee waggle dance: can we follow the steps? Trends in Ecology and Evolution, 24, 242–247.

45. Beekman, M., Gilchrist, A. L., Duncan, M., & Sumpter, D. J. T. (2007). What makes a honeybee scout? Behavioral Ecology and Sociobiology, 61, 985–995.

46. Hasenjager, M. J., Hoppitt, W., & Leadbeater, E. (2021). Data from: Honeybee dance-followers respond similarly to dances regardless of their spatial information content. Dryad Digital Repository. https://doi.org/10.5061/dryad.8kprr4xn8

