## Supplemental methods, figures, and tables for "Honeybee dance-followers respond similarly to dances regardless of their spatial information content"

### Supplementary methods

#### *Network-based diffusion analysis (NBDA)*

To compare the relative importance of waggle dance information in aiding discovery of the target feeder at different foraging distances, we carried out a network-based diffusion analysis (NBDA). NBDA estimates the extent to which interacting with knowledgeable individuals (represented using network data) accelerates an individual's learning of a novel behavioural pattern [1,2], taken here as discovery of the target feeder's location. The basic NBDA model can be formulated as:

$$\lambda_i(t) = \lambda_0(t)(1 - z_i(t))\left(s\sum_{j=1}^{N} a_{ij}z_j(t) + 1\right)$$

Eqn. 1

where $\lambda_i(t)$ is the rate at which individual *i* learns the novel behaviour as a function of time, $\lambda_0(t)$ is the baseline rate function that describes how learning rates change over time, *s* is the estimated rate of social transmission per unit of network connection with informed individuals, *N* is the number of individuals in the group or population, $a_{ij}$ is the network connection strength from individual *j* to *i*, and $z_j(t)$ indicates the status of individual *j* at time *t* (1 = informed, 0 = naïve). This basic model can be extended in several ways, such as allowing networks to change over time or incorporating additional variables that can modify asocial and/or social learning rates [2,3].

Here, we used an NBDA variant known as order-of-acquisition diffusion analysis (OADA), which takes as data the order in which recruits discovered the target feeder within each trial [2]. As OADA makes no assumptions about how learning rates are expected to change across time—simply assuming that all individuals within a trial share the same baseline rate function—it is well-suited for situations where the rate of learning is thought to fluctuate unpredictably [3]. Due to variation in weather conditions within and across trials (e.g. precipitation, strong winds), we had reason to suspect this was true for our data.

NBDA estimates the rate of social transmission, *s*, per unit of social network connection with informed individuals, relative to the baseline learning rate (Eqn. 1). This means that all else being equal, individuals that share more or stronger connections with informed group mates are expected to learn sooner than more weakly connected individuals. From our records of dance-following interactions, we constructed two dance-following networks: one in which edge weights equalled the number of waggle runs followed and another in which edges were weighted by the total duration (sec) of dance-following interactions. To capture the temporal ordering of dance-following interactions, we used dynamic networks that updated when the next individual to discover the target feeder was last observed in hive. Occasionally, a recruit left the hive before another recruit, but discovered the target feeder after it; in these instances, to avoid having networks "rewind", the update time for the latter was used for the former. To evaluate the relative importance of dance-based transmission across our distance treatments, we compared the fit of models in which $s$ was either estimated separately for short- and long-distance trials ($s_{Short} \neq s_{Long}$) or in which $s$ was constrained to be equal across these treatments ($s_{Short} = s_{Long}$).

We included two additional network types in our analysis in order to evaluate the possibility that neither dance network accurately reflected the pathways of transmission. In these networks, a binary connection (strength of 1) was formed between each informed bee and potential recruit either when they both co-occurred in the hive (co-occurrence network) or when the latter visited the hive for the first time since the former became informed (first-visit network). The former represents any opportunities for direct information transfer from informed individuals, whereas the latter represents opportunities for indirect transfer (e.g. receiving scented sucrose via trophallaxis from a third-party).

We further included in the NBDA the number of revisits to the recruit feeder (continuous; standardised within trials) to evaluate its effect on individual and/or social learning rates. It is not possible in an OADA to include variables that do not vary among individuals within a trial, meaning we could not include explicitly include effects of colony or trial [2].

All candidate models were run for the following combinations of parameters: network type (waggle runs followed/duration of dance-following/co-occurrence/first visit), social transmission parameterisation ($s_{Short} = s_{Long}$/$s_{Short} \neq s_{Long}$/$s_{Short} = 0$/$s_{Long} = 0$), and the effect of revisits to the recruit feeder (asocial effect/social effect/asocial and social effect/no effect). A pair of asocial models (i.e. $s_{Short} = s_{Long} = 0$) were also included, either with or without an asocial effect of revisits to the recruit feeder on the rate of target feeder discovery. We used AICc to compare support for all candidate models. To limit retention of overly complex models, we removed a model from the model set if it was a more complex version of a model that received greater support (i.e. the latter was nested within the former; [4]). Profile likelihood techniques were used to obtain 95% confidence intervals due to frequent asymmetry in the uncertainty associated with parameters [5]. The NBDA was carried out using the NBDA package v. 0.9.4 in R v. 4.0.3 [6,7].

**Table S1.** Weather conditions during trials

| Colony | TF Distance | TF Heading | Scent | Date | Temp. (°C) | Wind Direction | Wind Speed | Cloud cover |
|---|---|---|---|---|---|---|---|---|
| A | 100 m | SW | Eucalyptus | Jul. 28 | 21.5 | SW | 17 | 3 |
| A | 500 m | WSW | Bergamot | Aug. 18 | 20.8 | WSW | 11 | 7 |
| B | 100 m | ESE | Tea Tree | Sept. 4 | 18.8 | NNE | 7 | 8 |
| B | 500 m | ESE | Cypress | Aug. 29 | 18.3 | NW | 7 | 5 |
| C | 100 m | SW | Palma rosa | Sept. 20 | 19.6 | SSW | 15 | 8 |
| C | 500 m | WSW | Sweet orange | Sept. 13 | 19.5 | W | 7 | 0 |

Target feeder (TF) heading is given relative to the hive. Trials took place during 2018 and began between 0934 and 0952 UTC. Approximate daily weather conditions (1200 UTC) obtained from the Met Office using data from the nearest station (Heathrow; located about 9 km from Royal Holloway, University of London). Wind direction indicates the direction from which the wind is blowing from. Wind speed is given in knots. Cloud cover is given in Oktas, ranging from 0 (clear) to 8 (overcast). Some precipitation occurred for about 30 min during the 100 m trial for Colony C, though foragers continued to visit the TF feeder throughout this time.

#### *Individual-level analyses*

For the analysis of dancer and follower behaviour, we employed an information-theoretic approach in which a global model of interest was specified for each response variable. We then evaluated the relative support for all candidate models nested within each global model as described in the main text. In tables S2 and S3, we present the structure of the global models used in the analysis of follower and dancer behaviour respectively. For each model, we indicate the response variable, the number of observations, and the fixed and random effect structures. We also include additional information for each analysis—e.g. incorporation of variance structures, removal of outliers, how response variables were calculated. Data and code to reproduce all analyses can be found at [URL to be added upon manuscript acceptance].

**Table S2.** Global models for analysis of follower behaviour

| Model 1: Probability of searching for the target feeder; binomial GLMM; N = 1048 observations | | |
|---|---|---|
| Fixed effects | Random intercepts | Additional information |
| TF distance (100 m/500 m)<br><br>Recruited (Y/N)<br><br>Hive visit<br><br>TF distance * Recruited<br><br>TF distance * Hive visit<br><br>Recruited * Hive visit<br><br>Colony (A/B/C) | Trial ID<br>(6 trials)<br><br>Forager ID<br>(158 foragers) | Prior to analysis, observations were removed in which search behaviour could not be assigned due to termination of the trial. |
| Model 2: Waggle runs followed prior to departure; zero-inflated negative binomial GLMM; N = 924 observations | | |
| Fixed effects | Random intercepts | Additional information |
| *Count model:*<br>TF distance (100 m/500 m)<br><br>Search (React./Search)<br><br>Recruited (Y/N)<br><br>Hive visit<br><br>TF distance * Search<br><br>TF distance * Recruited<br><br>TF distance * Hive visit<br><br>Search * Recruited<br><br>Search * Hive visit<br><br>Recruited * Hive visit<br><br>Colony (A/B/C)<br><br>*Zero-inflated model:*<br>Hive visit | Trial ID<br>(6 trials)<br><br>Forager ID<br>(158 foragers) | Prior to analysis, observations were removed in which either no dances for the target feeder occurred in the hive or in which search behaviour could not be assigned due to termination of the trial. |

**Table S2 (cont).** Global models for analysis of follower behaviour

| Model 3: Duration (min) of hive absences; LMM; N = 928 observations | | |
|---|---|---|
| Fixed effects | Random intercepts | Additional information |
| TF distance (100 m/500 m)<br>Search (React./Search)<br>Hive visit<br>TF distance * Search<br>TF distance * Hive visit<br>Search * Hive visit<br>Colony (A/B/C) | Trial ID<br>(6 trials)<br>Forager ID<br>(157 foragers) | Response was log-transformed to ensure normally distributed residuals.<br>Incorporated variance structure that allowed for heterogeneous residual spread across reactivation vs. search trips. |
| **Model 4: Mean duration (min) of searches vs. collection trips; LMM; N = 275 observations** | | |
| Fixed effects | Random intercepts | Additional information |
| TF distance (100 m/500 m)<br>Trip type (Search/Collect)<br>TF distance * Trip type<br>Colony (A/B/C) | Trial ID<br>(6 trials)<br>Forager ID<br>(219 foragers) | Data included the mean duration of either unsuccessful searches for the target feeder made by potential recruits or collection trips to the target feeder (minus time spent at the feeder).<br>Response was log-transformed to ensure normally distributed residuals.<br>Incorporated variance structure that allowed for heterogeneous residual spread across collection vs. search trips. |
| **Model 5: Unsuccessful searchers before TF discovery; Poisson GLMM; N = 74 observations** | | |
| Fixed effects | Random intercepts | Additional information |
| TF distance (100 m/500 m)<br>Colony (A/B/C) | Trial ID<br>(6 trials) | Only includes individuals that were successfully recruited to the target feeder. |

**Table S3.** Global models for analysis of dancer behaviour

| | | |
|---|---|---|
| Model 1: Probability of dancing on each hive visit; binomial GLMM; N = 1868 observations | | |
| Fixed effects | Random intercepts | |
| TF distance (100 m/500 m)<br>Recruited (Y/N)<br>Time of visit<br>TF distance * Time of visit<br>TF distance * Recruited<br>Colony (A/B/C) | Trial ID<br>(6 trials)<br>Forager ID<br>(140 foragers) | |
| Model 2: Mean waggle runs produced; LMM; N = 109 observations | | |
| Fixed effects | Random intercepts | Additional information |
| TF distance (100 m/500 m)<br>Recruited (Y/N)<br>TF distance * Recruited<br>Colony (A/B/C) | Trial ID<br>(6 trials) | Response variable was calculated across hive visits during which an individual danced.<br>Incorporated variance structure that allowed for heterogeneous residual spread across TF distances, forager history (i.e. trained vs. recruited to the target feeder), and colonies. |
| Model 3: Frequency of hive visits (visits $hr^{-1}$); LMM; N = 141 observations | | |
| Fixed effects | Random intercepts | Additional information |
| TF distance (100 m/500 m)<br>Recruited (Y/N)<br>TF distance * Recruited<br>Colony (A/B/C) | Trial ID<br>(6 trials) | Visitation rates measured following first arrival at target feeder<br>Incorporated variance structure that allowed for heterogeneous residual spread across TF distances. |
| Model 4: Mean hive visit duration (min); LMM; N = 139 observations | | |
| Fixed effects | Random intercepts | Additional information |
| TF distance (100 m/500 m)<br>Recruited (Y/N)<br>TF distance * Recruited<br>Colony (A/B/C) | Trial ID<br>(6 trials) | Mean visit duration was calculated only for observed visits of known duration.<br>An observation with a mean visit duration of 31.1 min was identified as influential based on Cook's distance. Inclusion of this observation substantively altered conclusions drawn from the analysis and so was removed.<br>Incorporated variance structure that allowed for heterogeneous residual spread across TF distances, forager history (i.e. trained vs. recruited to the target feeder), and colonies. |

**Table S4.** Best-supported models from the network-based diffusion analysis

| Parameter | Estimate | 95% CI |
|---|---|---|
| Model 1 ($w_i = 0.908$) | | |
| ***S*** | **2.417 x 10$^7$** | **0.903, +∞** |
| **Number of revisits (social effect)** | **-0.535** | **-0.81, -0.275** |
| Model 2 ($w_i = 0.092$) | | |
| ***s*** | **1.212 x 10$^7$** | **0.364, +∞** |
| **Number of revisits (social effect)** | **-0.539** | **-0.815, -0.278** |

In both models, the rate of social transmission, $s$, was constrained to be equal between the short- and long-distance treatments (i.e. $s_{100m} = s_{500m}$). Connections in the dance-following network were either weighted by the number of waggle runs followed (Model 1) or the duration (sec) of dance-following interactions (Model 2). Out of the 74 recruitment events, Model 1 estimated dance-based transmission was responsible for 97.07 – 100% of events, whereas Model 2 estimated that dance-based transmission was responsible for 96.93 – 100% of events.

**Table S5.** Binomial GLMM of followers' probability of searching for the target feeder

| Parameter | MAE | USE | 95% CI |
|---|---|---|---|
| Intercept | 0.184 | 0.108 | -0.027, 0.396 |
| **Distance (500 m)** | **0.897** | **0.215** | **0.476, 1.318** |
| **Hive visit** | **2.697** | **0.229** | **2.247, 3.146** |
| **Recruited (Y)** | **0.891** | **0.207** | **0.486, 1.297** |
| **Distance * Hive visit** | **1.909** | **0.458** | **1.012, 2.806** |
| Distance * Recruited | 0.675 | 0.507 | -0.319, 1.67 |
| Hive visit * Recruited | 0.369 | 0.434 | -0.481, 1.219 |
| **Colony B** | **-1.369** | **0.253** | **-1.865, -0.873** |
| Colony C | -0.404 | 0.248 | -0.889, 0.081 |

Model-averaged estimates (MAE) and unconditional standard errors (USE) derived from the 3 best-supported models ($\sum w_i = 0.99$). Trial ID and forager ID were included as random intercept terms in all models. *N* = 1048 observations.

**Table S6.** Model set used to derive parameter estimates presented in table S5

| Model | Parameters | df | log *L* | AICc | ΔAICc | $w_i$ |
|---|---|---|---|---|---|---|
| 1 | Dist + Vis + Rec + Dist * Vis + Dist * Rec + Vis * Rec + Col | 11 | -596.791 | 1215.8 | 0 | 0.539 |
| 2 | Dist + Vis + Rec + Dist * Vis + Dist * Rec + Col | 10 | -598.505 | 1217.2 | 1.39 | 0.27 |
| 3 | Dist + Vis + Rec + Dist * Vis + Col | 9 | -599.922 | 1218.0 | 2.18 | 0.181 |

Dist = Target feeder distance (100 m / 500 m); Vis = Hive visit; Rec = Recruited (Trained / Recruited); Col = Colony ID.

**Table S7.** Zero-inflated negative binomial GLMM of the number of waggle runs followed per hive visit by followers

| Parameter | MAE | USE | 95% CI |
|---|---|---|---|
| *Count model* | | | |
| Intercept | 2.024 | 0.114 | 1.802, 2.247 |
| Distance (500 m) | 0.222 | 0.219 | -0.208, 0.651 |
| **Hive visit** | **0.493** | **0.068** | **0.36, 0.626** |
| Recruited (Y) | 0.049 | 0.065 | -0.078, 0.176 |
| **Searching (Target)** | **0.752** | **0.06** | **0.634, 0.87** |
| Distance * Searching | 0.177 | 0.148 | -0.114, 0.467 |
| Distance * Hive Visit | -0.024 | 0.074 | -0.168, 0.121 |
| Hive Visit * Recruited | 0.186 | 0.138 | -0.084, 0.457 |
| **Hive Visit * Searching** | **-0.529** | **0.13** | **-0.784, -0.274** |
| Recruited * Searching | -0.024 | 0.074 | -0.168, 0.12 |
| Colony B | 0.158 | 0.237 | -0.305, 0.622 |
| Colony C | -0.013 | 0.126 | -0.26, 0.235 |
| *Zero-inflated model* | | | |
| Intercept | -3.493 | 0.507 | -4.487, -2.499 |
| **Hive Visit** | **-3.699** | **1.145** | **-5.944, -1.455** |

Model-averaged estimates (MAE) and unconditional standard errors (USE) derived from the 8 best-supported models ($\sum w_i = 0.971$). Trial ID and forager ID were included as random intercept terms in all models. *N* = 924 observations.

**Table S8.** Model set used to derive parameter estimates presented in table S7

| Model | Parameters | Df | log *L* | AICc | ΔAICc | $w_i$ |
|---|---|---|---|---|---|---|
| 1 | Dist + Vis + Rec + Srch + Dist * Srch + Vis * Rec + Vis * Srch + Col + ZI(Vis) | 15 | -2673.92 | 5378.4 | 0 | 0.246 |
| 2 | Dist + Vis + Rec + Srch + Dist * Srch + Vis * Rec + Vis * Srch + ZI(Vis) | 13 | -2676.035 | 5378.5 | 0.1 | 0.234 |
| 3 | Vis + Rec + Srch + Vis * Rec + Vis * Srch + Rec * Srch + ZI(Vis) | 12 | -2677.614 | 5379.6 | 1.2 | 0.135 |
| 4 | Vis + Rec + Srch + Rec * Vis + Vis * Srch + ZI(Vis) | 11 | -2678.794 | 5379.9 | 1.51 | 0.116 |
| 5 | Dist + Vis + Srch + Dist * Srch + Dist * Vis + Vis * Srch + Col + ZI(Vis) | 14 | -2676.26 | 5381.0 | 2.61 | 0.067 |
| 6 | Dist + Vis + Srch + Dist * Srch + Dist * Vis + Vis * Srch + ZI(Vis) | 12 | -2678.375 | 5381.1 | 2.72 | 0.063 |
| 7 | Dist + Vis + Srch + Dist * Srch + Vis * Srch + Col + ZI(Vis) | 13 | -2677.42 | 5381.2 | 2.87 | 0.059 |
| 8 | Dist + Vis + Srch + Dist * Srch + Vis * Srch + ZI(Vis) | 11 | -2679.611 | 5381.5 | 3.14 | 0.051 |

Dist = Target feeder distance (100 m / 500 m); Vis = Hive visit; Rec = Recruited (Trained / Recruited); Srch = Searching (Reactivation / Searching); Col = Colony ID; ZI() = Term included in zero-inflated model.

**Table S9.** Linear mixed effects model of duration of hive absences (log-transformed) for potential recruits

| Parameter | MAE | USE | 95% CI |
|---|---|---|---|
| Intercept | 1.828 | 0.077 | 1.676, 1.979 |
| Distance (500 m) | 0.346 | 0.217 | -0.079, 0.771 |
| **Hive visit** | **0.192** | **0.031** | **0.132, 0.251** |
| Recruited (Y) | 0.047 | 0.051 | -0.054, 0.147 |
| Searching (Target) | -0.018 | 0.03 | -0.077, 0.042 |
| **Distance * Searching** | **0.6** | **0.063** | **0.476, 0.724** |
| Hive visit * Recruited | 0.022 | 0.054 | -0.083, 0.127 |
| **Hive visit * Searching** | **0.226** | **0.06** | **0.11, 0.343** |
| Recruited * Searching | 0.118 | 0.089 | -0.055, 0.292 |

Model-averaged estimates (MAE) and unconditional standard errors (USE) derived from the 3 best-supported models ($\sum w_i = 0.99$). Trial ID and forager ID were included as random intercept terms. A variance structure was incorporated that allowed residual spread to vary across reactivation vs. search trips. *N* = 928 observations.

**Table S10.** Model set used to derive parameter estimates presented in table S9

| Model | Parameters | df | log *L* | AICc | ΔAICc | $w_i$ |
|---|---|---|---|---|---|---|
| 1 | Dist + Vis + Rec + Srch + Dist * Srch + Rec * Srch + Vis * Srch | 12 | -506.662 | 1006.1 | 0 | 0.725 |
| 2 | Dist + Vis + Rec + Srch + Dist * Srch + Vis * Rec + Vis * Srch | 12 | -508.06 | 1009.0 | 2.95 | 0.174 |
| 3 | Dist + Vis + Srch + Dist * Srch + Vis * Srch | 10 | -506.646 | 1010.1 | 3.98 | 0.099 |

Dist = Target feeder distance (100 m / 500 m); Vis = Hive visit; Rec = Recruited (Trained / Recruited); Srch = Searching (Reactivation / Searching).

**Table S11.** Linear mixed effects model of mean duration of search vs. collection trips (log-transformed) for potential recruits and informed individuals respectively

| Parameter | Estimate | SE | 95% CI |
|---|---|---|---|
| Intercept | 1.559 | 0.023 | 1.514, 1.605 |
| **Distance (500 m)** | **0.627** | **0.03** | **0.495, 0.758** |
| **Trip type (Collection)** | **-0.598** | **0.046** | **-0.69, -0.507** |
| **Colony B** | **-0.267** | **0.037** | **-0.427, -0.108** |
| **Colony C** | **-0.189** | **0.038** | **-0.354, -0.025** |

Estimates derived from the best-supported model ($w_i = 0.999$). Trial ID and forager ID were included as random intercept terms. A variance structure was incorporated that allowed residual spread to vary across search vs. collection trips. *N* = 275 observations.

**Table S12.** Poisson GLMM of the number of unsuccessful searches made by recruits prior to successfully discovering the target feeder

| Parameter | MAE | USE | 95% CI |
|---|---|---|---|
| Intercept | 0.86 | 0.086 | 0.692, 1.027 |
| Distance (500 m) | -0.243 | 0.218 | -0.669, 0.183 |
| Colony B | 0.228 | 0.23 | -0.224, 0.679 |
| Colony C | 0.387 | 0.315 | -0.231, 1.005 |

Model-averaged estimates (MAE) and unconditional standard errors (USE) derived from the 2 best-supported models ($\sum w_i > 0.99$). Trial ID was included as a random intercept term. *N* = 74 observations.

**Table S13.** Model set used to derive parameter estimates presented in table S12

| Model | Parameters | df | log *L* | AICc | ΔAICc | $w_i$ |
|---|---|---|---|---|---|---|
| 1 | Dist + Col | 5 | -139.289 | 289.5 | 0 | 0.695 |
| 2 | -- | 2 | -143.469 | 291.1 | 1.65 | 0.305 |

Dist = Target feeder distance (100 m / 500 m); Col = Colony ID.

**Table S14.** Linear mixed effects model of the number of hive visits per hr by employed foragers

| Parameter | Estimate | SE | 95% CI |
|---|---|---|---|
| Intercept | 8.948 | 0.185 | 8.581, 9.314 |
| **Distance (500 m)** | **-3.581** | **0.365** | **-5.15, -2.011** |
| Colony B | 1.513 | 0.449 | -0.419, 3.445 |
| Colony C | 1.345 | 0.455 | -0.615, 3.305 |

Estimates derived from the best-supported model ($w_i = 0.95$). Trial ID was included as a random intercept term. A variance structure was incorporated that allowed residual spread to vary across TF distances. *N* = 141 observations.

**Table S15.** Linear mixed effects model of mean duration (min) of hive visits by employed foragers

| Parameter | Estimate | SE | 95% CI |
|---|---|---|---|
| Intercept | 1.576 | 0.051 | 1.476, 1.676 |
| **Distance (500 m)** | **0.506** | **0.089** | **0.121, 0.891** |
| **Recruited (Y)** | **-0.293** | **0.074** | **-0.44, -0.146** |
| Colony B | -0.288 | 0.138 | -0.883, 0.306 |
| Colony C | -0.469 | 0.134 | -1.044, 0.106 |

Estimates derived from the best-supported model ($w_i = 0.97$). Trial ID was included as a random intercept term. A variance structure was incorporated that allowed residual spread to vary across TF distances, forager history, and colonies. *N* = 139 observations.

**Table S16.** Binomial generalized linear mixed effects model of the probability of an employed forager dancing per hive visit

| Parameter | Estimate | SE | 95% CI |
|---|---|---|---|
| Intercept | 0.58 | 0.38 | -0.166, 1.325 |
| **Time of visit (sec)** | **1.954** | **0.168** | **1.625, 2.284** |
| **Recruited (Y)** | **-2.589** | **0.391** | **-3.354, -1.824** |

Estimates derived from the best-supported model ($w_i > 0.999$). Trial ID and forager ID were included as random intercept terms. Time of visit indicates when a forager arrived in the hive, measured as time elapsed since the start of the trial. *N* = 1868 observations.

**Table S17.** Linear mixed effects model of mean number of waggle runs produced per hive visit in which a forager danced for the target feeder

| Parameter | MAE | USE | 95% CI |
|---|---|---|---|
| Intercept | 15.18 | 1.849 | 11.557, 18.803 |
| Distance (500 m) | 0.308 | 1.79 | -3.201, 3.816 |
| Recruited (Y) | -3.576 | 2.005 | -7.506, 0.354 |
| Colony B | 0.162 | 2.216 | -4.182, 4.506 |
| Colony C | -0.424 | 2.519 | -5.36, 4.512 |

Model-averaged estimates (MAE) and unconditional standard errors (USE) derived across all three candidate models. Trial ID was included as a random intercept term. A variance structure was incorporated that allowed residual spread to vary across TF distances, forager history, and colonies. $N$ = 109 observations.

**Table S18.** Model set used to derive parameter estimates presented in table S17

| Model | Parameters | df | log $L$ | AICc | ΔAICc | $w_i$ |
|---|---|---|---|---|---|---|
| 1 | Rec | 8 | -386.873 | 796.8 | 0 | 0.851 |
| 2 | Dist + Col | 10 | -382.981 | 801.5 | 4.7 | 0.081 |
| 3 | -- | 7 | -391.884 | 801.9 | 5.05 | 0.068 |

Dist = Target feeder distance (100 m / 500 m); Rec = Recruited (Trained / Recruited); Col = Colony ID.

**Potential cross trips by foragers**

An absence from the hive was labelled as a "reactivation" if the forager was observed at the empty recruit feeder; otherwise, it was assumed that the forager was searching for the target feeder (labelled as a "search" trip). However, it is possible that during reactivation events, foragers may also have attempted to locate the target feeder. For example, using a two-feeder set up similar to the one used here, Menzel and colleagues [8] employed harmonic radar to track the flight behaviour of honeybee recruits in the field. While searching for a novel, dance-indicated site, foragers were occasionally observed to make cross flights between the location indicated by the dance and their familiar, but no longer profitable, feeding site (and vice versa) without first returning to the hive.

The likelihood of performing cross flights was related to the metric distance between two sites in the field—i.e. no such flights were observed between sites separated by > 300 m. This finding suggests that cross flights were unlikely during our 500 m trials, as the recruit and target feeders were separated by ~545 m. However, it is plausible that cross flights occurred during 100 m trials, in which the recruit and target feeders were only separated by ~175 m. In other words, some trips labelled as reactivation events during the 100 m trials may have also involved searching for the target feeder.

In the absence of technologies enabling continuous tracking of honeybees in the field, it is impossible to know with certainty how foragers behaved when they were not present on the dance floor or at a feeder. However, several lines of evidence suggest that although cross flights may have occurred during 100 m, they were unlikely to be prevalent enough to greatly impact our findings.

First, reactivation events did not differ in duration between short- and long-distance trials (main text, figure 4a; Fig. S1; table S9), which would not be expected if reactivation trips during the former also commonly involved searches for the target feeder.

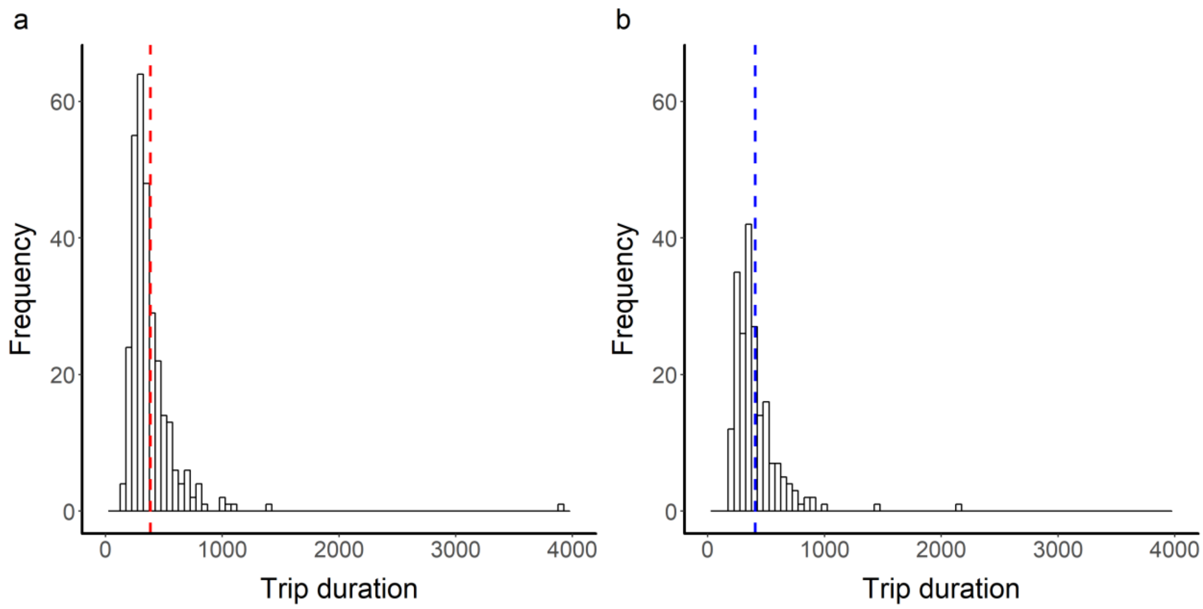


**Figure S1.** Duration of foraging trips labelled as reactivations for (a) 100 m and (b) 500 m trials. Dashed vertical lines indicate the mean duration.

The possibility remains that a forager might spend the majority of its time searching for the target feeder, while briefly checking the recruit feeder; such trips would still be classified solely as reactivation events under our criteria. If this were the case, then foragers would be expected to spend less time at the recruit feeder itself during 100 m trials, especially later in the trial when searching for the target feeder is more likely (table S5; main text, figure 2).

To evaluate this probability, we first constructed a linear mixed effects model (LMM) with the duration of time observed at the recruit feeder on each reactivation trip as a response variable and distance treatment, hive visit, the number of waggle runs followed during the preceding hive visit, and all two-way interactions involving distance as fixed effects. Trial and forager IDs were included as random effects and colony ID as a fixed effect. We then evaluated the support for all candidate models nested within this global model as described in the main text. We found that as trials progressed, foragers tended to spend less time at the recruit feeder (hive visit: MAE ± UCSE = -1.77 ± 0.42; table S19). However, there was no evidence that foragers in 500 m trials spent more time at the recruit feeder than foragers in 100 m trials (target feeder distance (500 m): MAE ± UCSE = -0.1 ± 0.85; table S19).

**Table S19.** Linear mixed effects model of duration of time (sec) observed at recruit feeder on each reactivation trip (square-root transformation applied)

| Parameter | MAE | USE | 95% CI |
|---|---|---|---|
| Intercept | 9.433 | 0.241 | 8.96, 9.906 |
| Distance (500 m) | -0.096 | 0.848 | -1.758, 1.566 |
| **Hive visit** | **-1.768** | **0.423** | **-2.596, -0.939** |
| **Colony B** | **3.485** | **1.276** | **0.984, 5.986** |
| Colony C | 0.785 | 1.268 | -1.7, 3.271 |
| Distance * Hive visit | -1.311 | 1.192 | -3.647, 1.025 |

Model-averaged estimates (MAE) and unconditional standard errors (USE) derived from the 2 best-supported models ($\sum w_i = 0.997$). Trial ID and forager ID were included as random intercept terms; colony ID was included as a fixed effect. *N* = 529 observations.

If reactivation events during 100 m trials frequently involved searches for the target feeder, we would also expect increased investment in following waggle dances by foragers prior to these trips. To evaluate this possibility, we first constructed a zero-inflated negative binomial GLMM with the number of waggle runs followed prior to each reactivation trip as the response variable and target feeder distance, hive visit, and their interaction as predictors in the count model. The zero-inflated model included an effect of hive visit. Trial and forager IDs were included as random effects and colony ID as a fixed effect. We then evaluated all subsets of this global model as described in the main text. As trials progressed, individuals tended to follow more waggle runs before departing the hive, even when considering only reactivation trips (hive visit: count model: estimate ± SE = 0.89 ± 0.14 (95% CI: 0.62, 1.15); table S20). However, there was no evidence that foragers in short-distance trials followed more dances prior to reactivation than those in long-distance trials (i.e. the top-ranked model, $w_i > 0.99$, did not include an effect of distance).

We also note that in none of the 74 successful searches did recruits first visit the recruit feeder prior to locating the target feeder without an intermediary visit to the hive. Furthermore, once a recruit located the target feeder, in no instances did we observe that individual returning to the recruit feeder afterwards.

Taken together, these results suggest that although cross trips may have occurred during our experiments (resulting in some search events being labelled as reactivations), it appears unlikely that they were sufficiently prevalent to substantially alter our findings.

**Table S20.** Zero-inflated negative binomial GLMM of number of waggle runs followed prior to reactivation events

| Parameter | Estimate | SE | 95% CI |
|---|---|---|---|
| *Count model* | | | |
| Intercept | 1.506 | 0.15 | 1.212, 1.8 |
| **Hive visit** | **0.887** | **0.135** | **0.622, 1.152** |
| *Zero-inflated model* | | | |
| Intercept | -3.78 | 0.764 | -5.278, -2.282 |
| **Hive visit** | **-5.811** | **1.45** | **-8.652, -2.97** |

Estimates derived from the best-supported model ($w_i > 0.99$). Trial ID and forager ID were included as random intercept terms. *N* = 423 observations.

**Analysis of follower behaviour during successful searches**

For the 74 successful recruitment events, we investigated whether the number of waggle runs followed prior to departing the hive in search of the target feeder or the search duration varied across short- and long-distance trials.

**Table S21.** Negative binomial GLMM of the number of waggle runs followed per hive visit by followers prior to a successful search.

| Parameter | MAE | USE | 95% CI |
|---|---|---|---|
| Intercept | 2.57 | 0.05 | 2.472, 2.668 |
| Distance (500 m) | 0.179 | 0.154 | -0.122, 0.48 |
| Hive visit | -0.049 | 0.104 | -0.252, 0.154 |
| **Colony B** | **0.51** | **0.14** | **0.237, 0.784** |
| Colony C | -0.088 | 0.163 | -0.407, 0.231 |

Model-averaged estimates (MAE) and unconditional standard errors (USE) derived from the 3 best-supported models ($\sum w_i = 0.994$). Trial ID was included as a random intercept terms in all models. *N* = 74 observations.

**Table S22.** Linear mixed effects model of duration of successful searches (log-transformed)

| Parameter | Estimate | SE | 95% CI |
|---|---|---|---|
| Intercept | 1.389 | 0.06 | 1.273, 1.505 |
| **Distance (500 m)** | **0.523** | **0.134** | **0.264, 0.782** |
| **Colony B** | **-0.723** | **0.152** | **-1.016, -0.431** |
| Colony C | -0.174 | 0.167 | -0.496, 0.148 |

Estimates and standard errors derived from the best-supported model ($\sum w_i = 0.983$). Trial ID was included as a random intercept term. *N* = 74 observations.

**Supplementary references**


1. Franz, M., & Nunn, C. L. (2009). Network-based diffusion analysis: a new method for detecting social learning. *Proceedings of the Royal Society B*, *276*, 1829-1836.

2. Hoppitt, W., Boogert, N. J., & Laland, K. N. (2010). Detecting social transmission in networks. *Journal of Theoretical Biology*, *263*, 544-555.

3. Hasenjager, M. J., Leadbeater, E., & Hoppitt, W. (2020). Detecting and quantifying social transmission using network-based diffusion analysis. *Journal of Animal Ecology*. doi: 10.1111/1365-2656.13307.

4. Richards, S. A. (2008). Dealing with overdispersed count data in applied ecology. *Journal of Applied Ecology*, *45*, 218-227.

5. Morgan, B. J. T. (2009). *Applied stochastic modelling* (2nd ed.) Boca Raton, FL: Chapman & Hall/CRC Press.

6. R Core Team. (2020). R: a language and environment for statistical computing. R Foundation for Statistical Computing, Vienna, Austria. https://www.R-project.org/

7. Hoppitt, W., Photopoulou, T., Hasenjager, M., & Leadbeater, E. (2020). NBDA: a package for implementing network-based diffusion analysis. R package version 0.9.4. https://github.com/whoppitt/NBDA

8. Menzel, R., Kirbach, A., Haass, W.-D., Fischer, B., Fuchs, J., Koblofsky, M., et al. (2011). A common frame of reference for learned and communicated vectors in honeybee navigation. *Current Biology*, *21*, 645-650.
